# Volumetric morphometry reveals mitotic spindle width as the best predictor of spindle scaling

**DOI:** 10.1101/2021.04.08.438956

**Authors:** Tobias Kletter, Sebastian Reusch, Nils Dempewolf, Christian Tischer, Simone Reber

## Abstract

The function of cellular structures at the mesoscale is dependent on their geometry and proportionality to cell size. The mitotic spindle is a good example why length and shape of intracellular organelles matter. Spindle length determines the distance over which chromosomes will segregate and spindle shape ensures bipolarity. While we still lack a systematic and quantitative understanding of subcellular morphometrics, new imaging techniques and volumetric data analysis promise novel insights into scaling relations across different species. Here, we introduce Spindle3D, an open-source plug-in that allows for the quantitative, unbiased, and automated analysis of 3D fluorescent data of spindles and chromatin. We systematically analyse different cell types, including somatic cells, stem cells and one-cell embryos across different phyla to derive volumetric relations of spindle, chromatin, and cell volume. Taken together, our data indicate that mitotic spindle width is a robust indicator of spindle volume, which correlates linearly with chromatin and cell volume both within single cell types and across metazoan phyla.

## Introduction

Size and shape in general are important biological features. Classically, morphometrics have been studied on the level of organisms, tissues, and cells. Now, the continuous improvement of imaging techniques and data analysis allows for the accurate measurement of organelle geometry at the µm-scale and thus enables the development of quantitative scaling laws at the mesoscale.

Anecdotal evidence suggested spindle length to scale with cell size. More recent studies, however, show that the nature of spindle scaling and size control is more complex. In different cells, spindle length spans over an order of magnitude and various molecular scaling mechanisms are likely to contribute to different degrees to cover the entire length regime (Rieckhoff et al., 2020). For example, the length of mitotic spindles increases with cell length in small cells, but in very large cells spindle length approaches an upper limit (Wuhr et al., 2008; Lacroix et al., 2018; Rieckhoff et al., 2020). More precisely, spindle length scales linearly with cytoplasmic volume (Hazel et al., 2013; Good et al., 2013). Further, spindle size needs to be coordinated with chromosome length, a fact that is established (Mora-Bermúdez et al., 2007; Lipp et al., 2007; Dinarina et al., 2009; Kieserman and Heald, 2011) but poorly understood. All the above observations point to an important, in some cases still open, question: What are the relevant morphometric measures to precisely formulate spindle scaling phenomena?

While most experimental studies still measure spindle length and cell diameter or collapse them into area information, theoretical arguments regularly use volumetric data (Good et al., 2013; Reber et al., 2013; Rieckhoff et al., 2020). However, most tools available for analysing spindle size and geometry only allow for 2D analysis (Crowder et al., 2015; Grenfell et al., 2016). Here, we show that spindle parameters differ significantly when measured in 2D as compared to 3D, they are thus error-prone and might lead to incorrect mechanistic conclusions. We argue that quantitative measurements from 3D datasets are essential to allow for accurate mechanistic interpretation and to derive conceptual scaling laws. Therefore, we use quantitative microscopy together with a newly developed analysis tool “Spindle3D” and the segmentation software Ilastik (Berg et al., 2019) to derive quantitative 3D morphometry data on spindle, chromatin, and cell volume. Our data imply that mitotic spindle width is a more robust predictor of spindle scaling than spindle length. Spindle3D is available free and open source as plug-in (https://sites.imagej.net/Spindle3D) in Fiji (Schindelin et al., 2012) and allows for the quantitative, unbiased, and automated analysis of 3D fluorescent data of spindles and chromatin. Our analysis of a variety of cell types and phyla proves the robustness of the Spindle3D plug-in, which thus can be broadly used by researchers to inform future biological and physical concepts of spindle scaling and size control across many experimental systems.

## Results

### 3D-analysis of fluorescent spindle and chromatin data allows for the accurate extraction of quantitative morphometric parameters

To extract quantitative parameters of spindle and chromatin morphometries, our plug-in requires a two-channel Z-series with fluorescent tubulin and chromatin labelings (Figure 1A and S1) as a minimum input, which allows for object detection and spindle axis localization (Material and Methods). Classically, spindle length is defined as the pole-to-pole distance. Consistently, we first define the spindle axis, along which spindle length is specified as the distance between the two spindle poles (Figure 1B). Next, spindle width is measured orthogonally to the spindle axis and reflects the diameter of the spindle ellipsoid at its equator. The volumes of all voxels within the segmented spindle mask add up to yield a spindle’s volume. As the orientation of the mitotic spindle can determine both, the relative size and the position of the daughter cells (as reviewed in McNally, 2013), we provide a measure for the spindle angle, which describes the tilt of the spindle axis relative to the reference plane. Chromatin can induce spindle assembly (Heald et al., 1996) and influences the spindle’s geometry (Dinarina et al., 2009). Hence, we use the intensity profile of the chromatin to measure the metaphase plate, which expands orthogonally to the determined spindle axis (Figure S1A). We define the metaphase plate width as the extent of chromatin along its shortest axis and the metaphase plate length as the mean chromatin diameter. Again, the volumes of all voxels within the segmented chromatin mask add up to yield chromatin volume. In some cell types, we consistently find the chromosomes aligned with a central opening in the metaphase plate, a phenomenon we termed chromatin dilation, which is measured using a radial intensity profile (Figure 1B and S1C).

**Figure 1:**
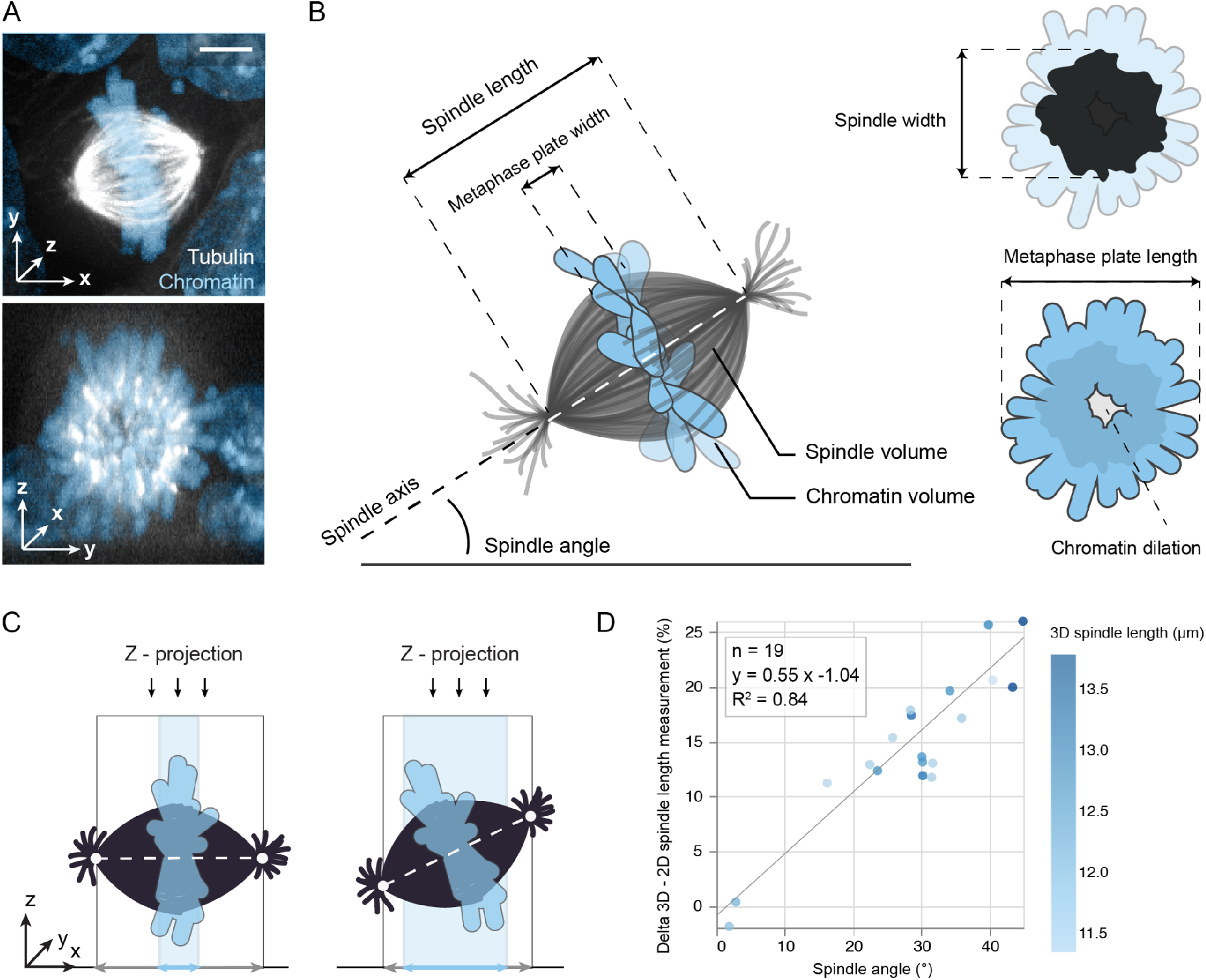
3D analysis of fluorescent spindle and chromatin data allows for the accurate extraction of morphometric parameters. (**A**) Top: projected micrograph of a mitotic mouse embryonic stem cell expressing tubulin-GFP (white), DNA stained with Hoechst (blue). Bottom: same image resliced to display the equatorial section of the spindle. Scale bar: 5 µm (**B**) Schematic of a mitotic spindle and its relevant morphometric parameters extracted by Spindle3D. Along the spindle axis, Spindle3D measures spindle length and metaphase plate width, and in the lateral direction spindle width and metaphase plate length. Chromatin dilation quantifies the central signal strength of the metaphase plate. (**C**) Morphometry on projected spindles distorts measurements, if spindle axes are tilted. (**D**) The relationship between the spindle angle and the percentaged discrepancy between the 2D (projected) and 3D spindle length quantification (n = 19). Circles represent individual spindles, colour is coded according to 3D spindle length. Line shows linear regression.

By projecting our 3D microscopic images into 2D planes (Figure 1C), we identified and quantified potential sources of error when spindle parameters are only analysed in two dimensions. Collapsing 3D into 2D information resulted in the distortion of morphometric parameters. The error is particularly evident when spindles are not perfectly parallel to the substrate but tilted in Z (Figure 1D). As a consequence, axial extents such as spindle length and metaphase plate width will be under- or overestimated, respectively. Even moderate spindle angles of 25° produce measurement errors of approximately 12.5% (Figure 1D). Furthermore, because spindle morphologies are not always perfectly symmetrical, information on spindle width and metaphase plate length are lost in 2D projections. Taken together, we recommend the acquisition and analysis of 3D datasets, which is essential to derive accurate quantitative measurements, in particular if they shall inform mathematical models.

### Spindle3D robustly derives morphometric parameters across a variety of cell types and phyla

To allow for an automated and quantitative analysis of spindle and chromatin parameters, we developed a 3D morphometric analysis workflow (Figure S1). In order to demonstrate its applicability and robustness, we subjected confocal images of live metaphase spindles from wild-type HEK293 cells (*Homo sapiens*), a HeLa Kyoto cell line (*H. sapiens*), Ptk_2_ cells (*Potorous tridactylis*), bovine zygotes (*Bos taurus*), and murine embryonic stem cells (mESCs, *Mus musculus*) to our 3D morphometric analysis (Figure 2A). The workflow produced segmented outputs with spindle axis-aligned voxel coordinates (Figure 2B). Analysed spindle lengths ranged from 6.8 - 26.5 µm (Figure 2C). Together with mESCs, the two human cell lines (HEK293 and HeLa Kyoto) showed spindles of comparable lengths (11.5±1.2 µm (mean±standard deviation), 11.5±1.4 µm, 13.6±2.0 µm, respectively), contrasted by the considerably longer spindles of the bovine zygotes and Ptk_2_ cells with an average length of 16.9±2.9 µm and 16.4±2.9 µm, respectively. However, together with mESCs, Ptk_2_ cells displayed narrower spindles (Figure 2D), producing markedly increased spindle aspect ratios (Figure 2E), a shape descriptor defined as the ratio of spindle length and width. We found that small aspect ratios often coincided with flat spindle poles, while spindles with large aspect ratios had visibly pointed poles. Of the data sets tested, bovine zygotes harboured spindles with the largest volumes (Figure 2F) of 2069.4±661.1 µm^3^ (mean±standard deviation), consistent with spindles reaching an upper limit in early development (Wühr et al., 2008). In contrast, spindles in murine embryonic stem cells occupied only a fraction of this volume (408.8±101.6 µm^3^). Independent of cell type, the majority of spindles showed tilted angles (Figure 2G) that fall within a range of 0 - 77°, highlighting the importance of 3D analysis. Intriguingly, the volume and length of the metaphase plate (Figure 2H-I), did neither reflect cell-type specific genome sizes nor chromosome numbers (Figure S3A-D), hinting towards different levels of chromatin compaction (as reviewed in Levy and Heald, 2012). In addition, our analysis quantified various levels of chromatin dilation. Especially in mESCs spindles, but also in a large fraction of the HeLa Kyoto population, chromatin plates were frequently and considerably dilated (chromatin dilation > 0.5) (Figure 2J). Taken together, the 3D analysis workflow provided by our plug-in robustly revealed cell-type specific spindle and chromatin morphology across different cell lines.

**Figure 2:**
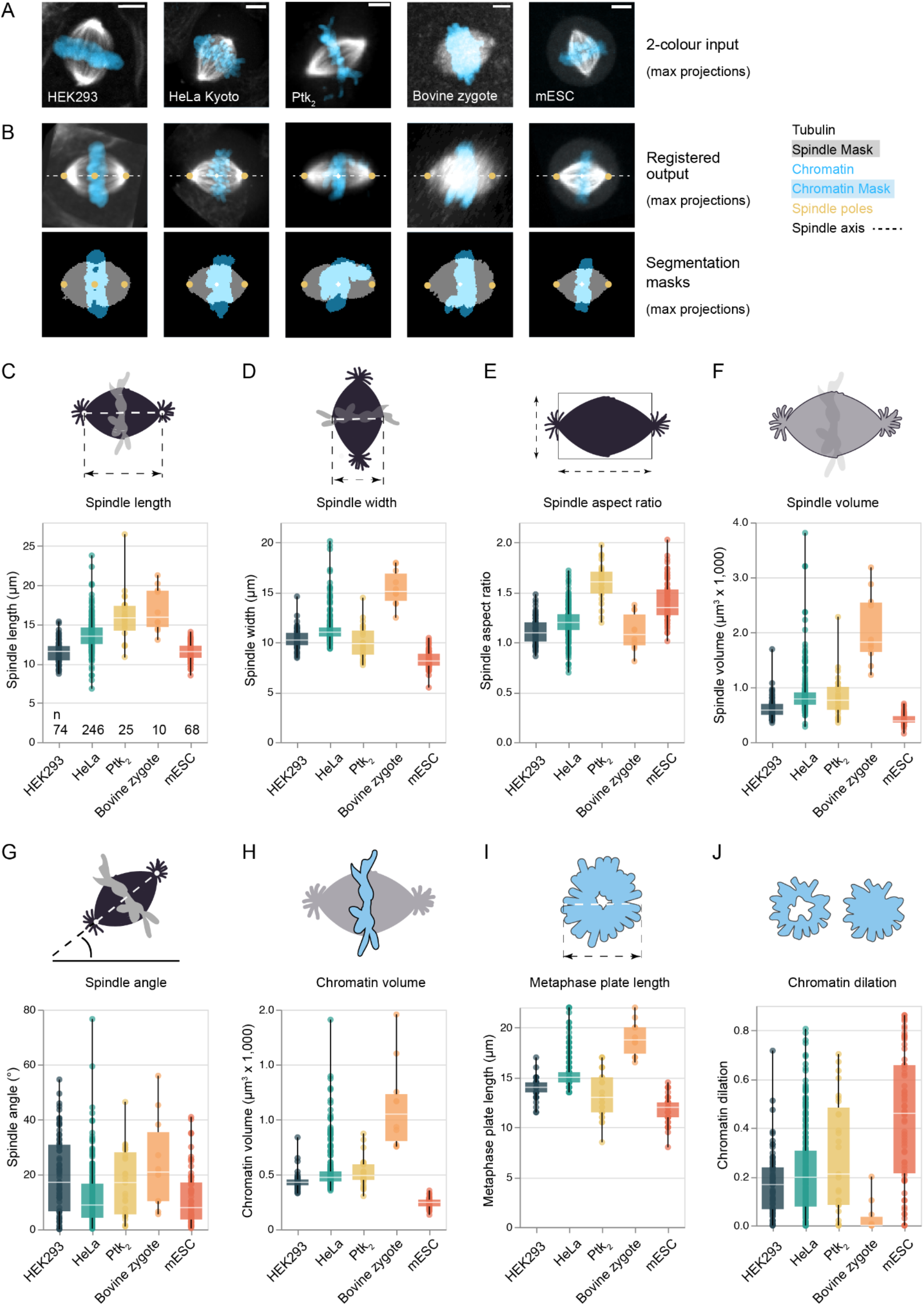
Spindle3D robustly derives morphometric parameters across a variety of cell types and phyla. (**A**) Representative live fluorescent 3D spindle data sets from different cells expressing labelled tubulin or microtubule-associated proteins or were treated with SiR-tubulin (white). Chromatin (blue) is visualized with Hoechst, SiR-DNA or H2B-mScarlet. Scale bar: 5 µm. (**B**) Automated axial registration and segmentation of 2-colour (tubulin gray scale, chromatin blue) input images as shown in (A). Spindle3D exports axially aligned output images containing segmentation masks and spindle pole localization for quality control. Quantification of (**C**) spindle length, (**D**) spindle width, (**E**) spindle aspect ratio, (**F**) spindle volume, (**G**) spindle angle, (**H**) chromatin volume, (**I**) metaphase plate length, and (**J**) chromatin dilation for all cell types. Circles are individual data points and represent a single spindle measurement. Boxes describe the interquartile range. Horizontal line in the box denotes median. Whiskers show minimum and maximum.

### Fixation and sample preparation alter spindle and chromatin morphology

In our explorations, we frequently observed a discrepancy in spindle sizes of live cells compared to cells that were fixed and mounted on cover slides. To systematically test the influence of fixation and mounting, we used HeLa Kyoto cells and mESCs stably expressing tubulin-GFP, allowing us to directly benchmark the mounted cells to their live counterparts (Figure 3A). Additionally, we measured cells that were chemically fixed, but not mounted. Using our plug-in, morphometric analysis revealed a marked decrease in spindle volumes in cells that were fixed and mounted in mounting media (Figure 3B). In addition, we frequently observed deformed spindle shapes in these samples (Figure S3) with shifted aspect ratios (Figure 3C). When samples were fixed but not mounted, spindle volumes were comparable to live spindles (Figure 3B). In these samples, as well as in the fixed + mounted samples, we observed a distorted volumetric relationship between spindle and chromatin (Figure 3D). Taken together, we show that sample preparation may induce artefacts in spindle and chromatin morphology and should be considered with great care. Importantly, as the introduced errors are not isotropic, such analyses may distort geometrical relationships and thus lead to error-prone scaling relations. Therefore, we recommend using live cells where possible, and for assays with fixed samples (e.g. immunofluorescence), we suggest refraining from mounting samples.

**Figure 3:**
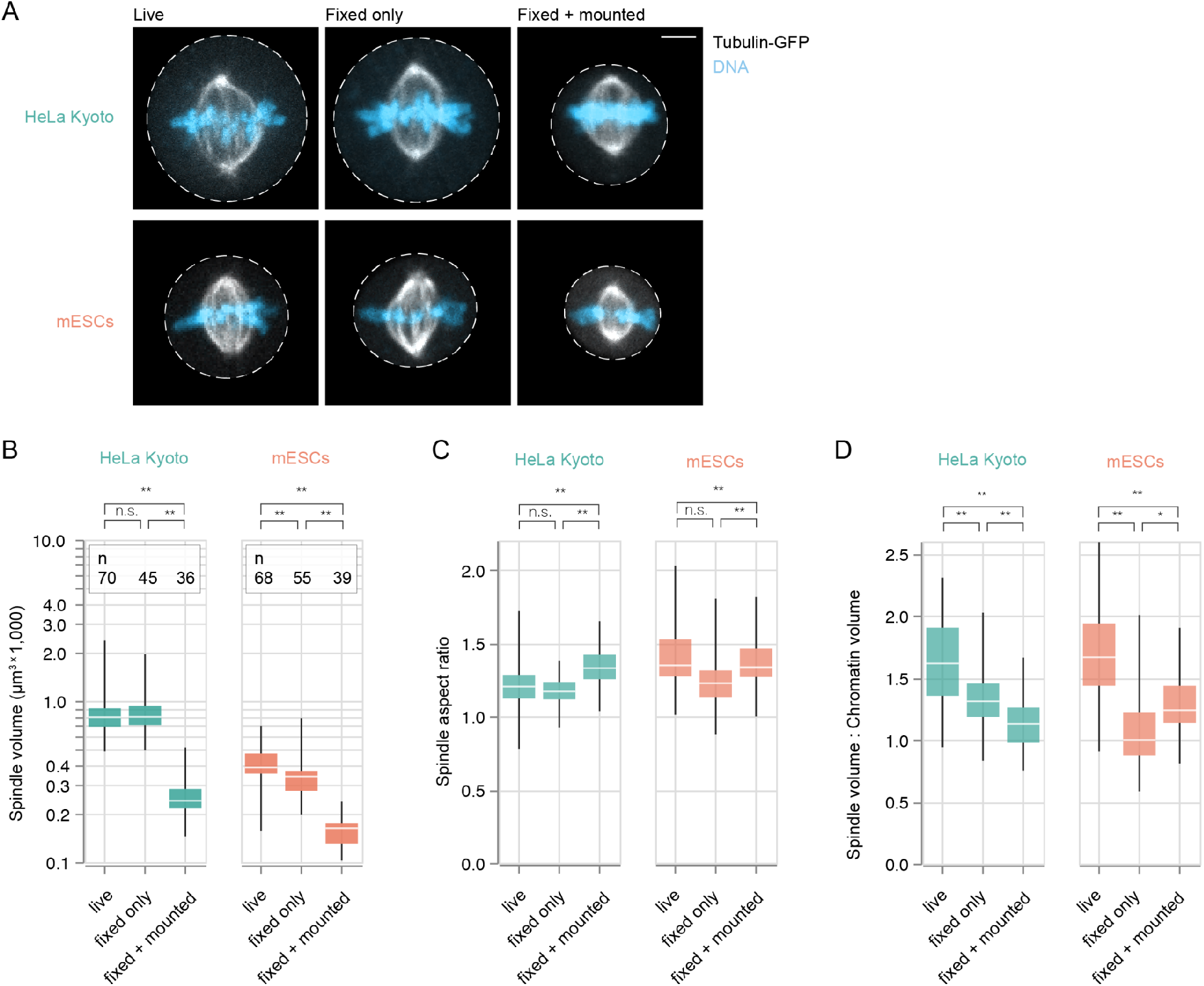
Fixation and sample preparation alter spindle and chromatin morphology. (**A**) Fluorescently-tagged tubulin allows for direct comparison of spindle morphology in live and fixed specimens. Left column shows representative mitotic cells (HeLa Kyoto and mESCs lines both stably expressing tubulin-GFP) when imaged live. Cells depicted in the central column were chemically fixed before imaging. Cells in the right column were fixed and embedded in mounting media. Tubulin-GFP signal is in white, DNA in blue. Dotted lines indicate cell boundaries. Scale bar: 5 µm. Fixation and sample preparation introduce artifacts to morphometric parameters such as (**B**) spindle volume and thus distort geometrical relationships among spindle measures such as (**C**) spindle aspect ratio and (**D**) the ratio of spindle volume to chromatin volume. Boxes denote interquartile range, horizontal lines show medians. Whiskers show minimum and maximum. P values from ANOVA with Tukey’s test as post-hoc analysis. * p < 0.05, ** p = 0.001, n.s: not significant (p > 0.05).

### Spindle width, rather than spindle length, reflects spindle volume

Several lines of evidence imply that there is a correlation between chromatin dimensions, spindle geometry, and steady-state microtubule polymer mass (as reviewed in Levy and Heald, 2012). From our analyses, we can now establish such simple scaling relations. We first explored the relationships between the length of the spindle and its volume (Figure 4A) and between the width of the spindle and its volume (Figure 4B). We found that, in both cases, the relationship can be described in terms of simple power laws. However, when calculating the spearman’s correlation coefficients r_s_, we found spindle width to correlate more strongly with spindle volume (r_s_ = 0.91, p = 2 × 10^-160^, Figure 4B) than we observed for the relationship between spindle length and volume (r_s_ = 0.77, p = 2 × 10^-83^, Figure 4A). This also held true when looking at the individual cell types (Figure 4A-B). As this was unexpected, we first verified manually that spindle length and width were represented reliably by our automated measurements (Figure S4A-D). Because the width of the spindle is in fact a three-dimensional parameter, we can measure spindle width as the average extent of the spindle mask projected along the spindle axis (Figure 4B) or as the maximum or minimum spindle width (Figure S4E-G). In either case, spindle width yielded strong correlations with spindle volume (Figure S4E-G). Chromatin has been shown to affect both spindle length and shape (Dinarina et al., 2009; Hara and Kimura, 2013). We thus plotted spindle volume as a function of chromatin volume (Figure 4C), which we find to correlate linearly (r_s_ = 0.72, p = 3 × 10^-69^). This is surprising because in embryonic systems, varying chromatin content only had a weak effect on spindle size while varying chromatin geometry influenced spindle assembly more drastically (Brown et al., 2007; Wühr et al., 2008; Dinarina et al., 2009). We therefore plotted spindle length and width against chromatin volume, and again found spindle width to correlate more strongly (r_s_ = 0.54, p = 2 × 10^-33^ for spindle length, Figure 4D and r_s_ = 0.73, p = 4 × 10^-72^ for spindle width, Figure 4E) with chromatin volume. Previous data implied that symmetric and thus functional spindles only self-organize around specific chromatin dimensions (Dinarina et al., 2009). Indeed, we find an almost perfect linear relation between spindle width and the length of the metaphase plate (r_s_ = 0.87, p = 1 × 10^-128^, Figure 4F) while spindle length and metaphase plate length only showed moderate dependencies (r_s_ = 0.44, p = 9 × 10^-22^). Taken together, we show that - contrary to common expectation - steady-state spindle width, rather than spindle length, is a reliable predictor of overall spindle volume.

**Figure 4:**
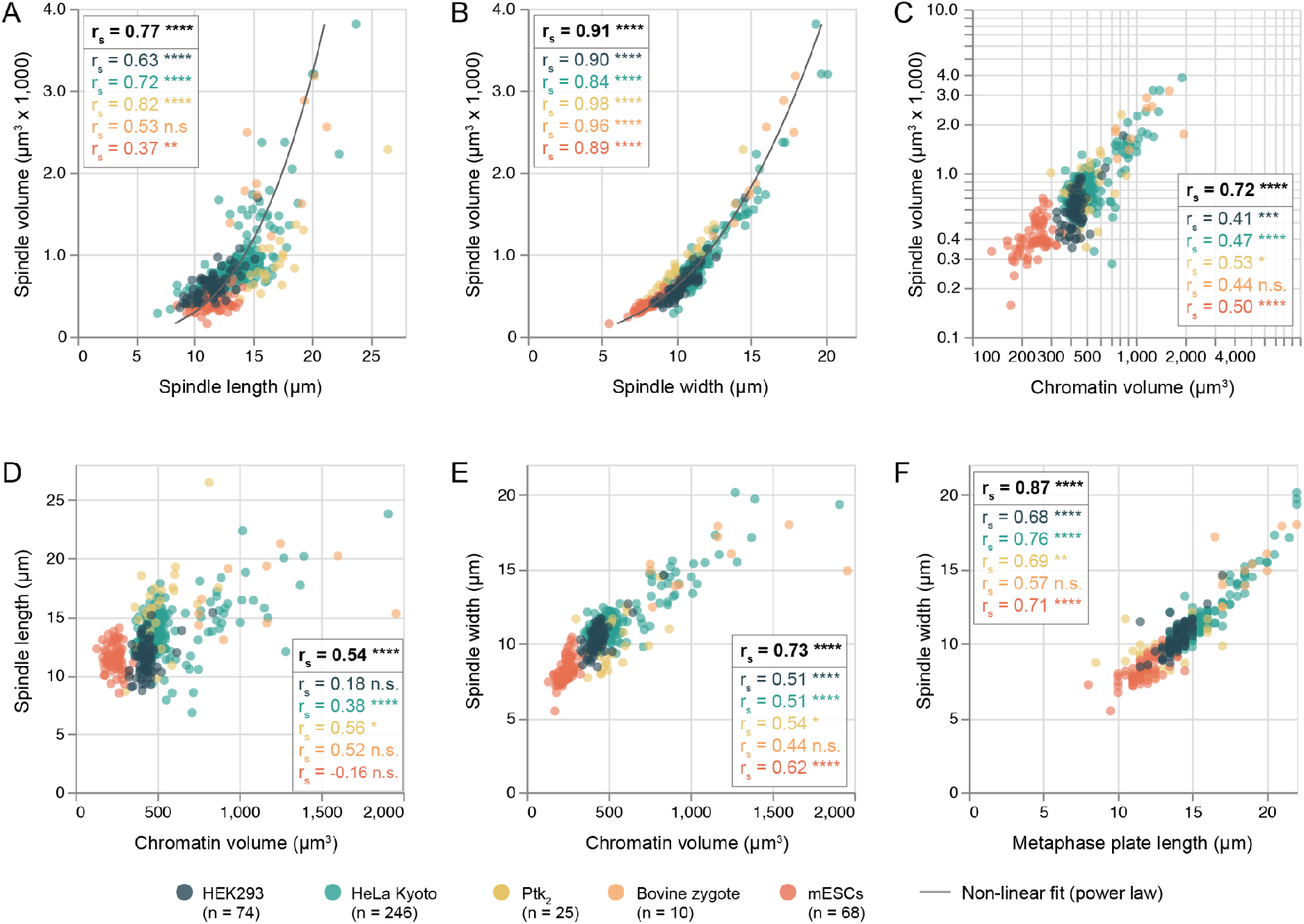
Spindle width, rather than spindle length, reflects spindle volume. Relationship between **(A)** spindle length and spindle volume and **(B)** spindle width and spindle volume in live spindles of five different cell types. **(C)** Volumetric relationship between chromatin and spindle. Relationship between **(D)** chromatin volume and spindle length and **(E)** chromatin volume and spindle width. **(F)** Relationship between metaphase plate length and spindle width. Circles represent individual cells. Individual data points in (A) and (B) fitted by a power law (f(x) = a x^k^, a: spindle length: 2.681 ± 0.535, exponent: 2.191 ± 0.073 (n = 425 cells); a: spindle width: 1.739 ± 0.001, exponent: 2.551 ± 0.000 (n = 425 cells)). r_s_: Spearman’s correlation coefficient, black coefficients show correlation for pooled data, coloured coefficients show cell-type resolved correlations. * p < 0.05, ** p < 0.005, *** p < 0.001, **** p < 0.0001, n.s.: not significant (p > 0.05).

### Spindle volume and chromatin volume scale linearly with cell volume

In many systems, cell or cytoplasmic volume is a major determinant of spindle size (Good et al., 2013; Hazel et al., 2013; Farhadifar et al., 2015; Wang et al., 2016; Lacroix et al., 2018; Rieckhoff et al., 2020). So, to reliably measure cell volume, we took advantage of the fact that mitotic cells expressing fluorescently-tagged tubulin show, next to the prominent spindle signal, distinctive fluorescence of soluble tubulin throughout the cytoplasm (Figure 5A). We thus used pixel classification-based 3D segmentation (Berg et al., 2019) of mitotic cells expressing fluorescent tubulin as a read-out of cell volume and cell sphericity to complement the morphometric data generated by our plug-in. Based on this, we trained random forest classifiers to distinguish and predict mitotic cell volumes (Figure 5B) and verified that this approach was as accurate as manual volumetric segmentations guided by cell-membrane labellings (Figure S5A-C). We live-imaged and analysed HeLa Kyoto, Ptk_2_ and mESCs expressing fluorescently-tagged tubulin. With an average cell volume of ≈ 6100 µm^3^ (HeLa Kyoto, Figure 5C), 7450 µm^3^ (Ptk_2_) and 2850 µm^3^ (mESCs), all cell types are expected to fall into the linear scaling regime (cell volume, V_c_ < 10^6^ µm^3^ (Rieckhoff et al., 2020), cell diameter, d_c_ < 140 µm, (Crowder et al., 2015)). In contrast to the other two cell lines, Ptk_2_ cells did not round up during mitosis (Figure 5D). Nevertheless, all three cell types displayed volumetric scaling between the cell and the respective spindle (r_s_ = 0.86, p = 3 × 10^-57^, Figure 5E). Interestingly, cells exerted a cell type-specific spindle size specification: While spindle volumes in both HeLa cells and mESCs occupied approximately 14% of total cell volumes, it was only 10% in Ptk_2_ cells (Figure 5F).

**Figure 5:**
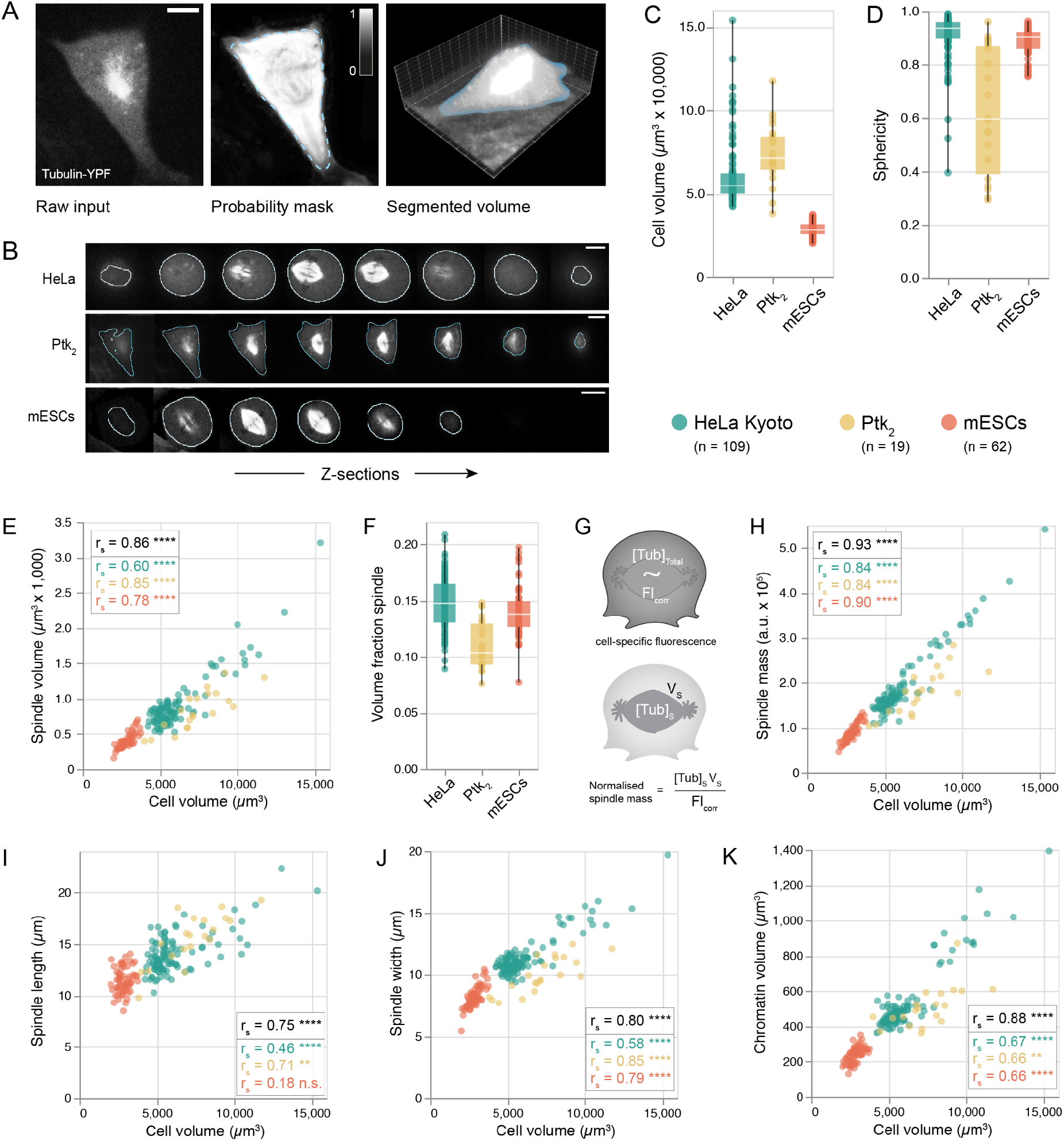
Spindle volume and chromatin volume scale linearly with cell volume. **(A)** Rationale for quantifying cell volumes via cytoplasmic tubulin fluorescence by pixel classification in the segmentation software Ilastik (Berg et al., 2019). Voxels of input micrographs (left) were converted to probabilities for mitotic cytoplasm (center). Probability masks were thresholded at 0.5 (dotted line) to produce the final volume mask (blue, right). Scale bar: 5 µm. **(B)** Z-series showing the cell boundaries (blue) as determined by pixel classification in three cell lines expressing fluorescent tubulin. Scale bars: 5 µm. **(C)** Distributions of cell volumes and **(D)** cell sphericity in three cell lines. **(E)** Bivariate relationships between cell volume and spindle volume and **(F)** distributions showing the fraction of cell volume occupied by the spindle. **(G)** Quantifying spindle mass as the polymer and free tubulin mass within the spindle volume, normalised by a cell-specific fluorescence correction factor proportional to the total fluorescent tubulin concentration. **(H)** Bivariate relationships between cell volume and spindle mass, **(I)** cell volume and spindle length and **(J)** cell volume and spindle width. **(K)** Volumetric relationship between spindle and chromatin. Circles reflect individual cells. r_s_: Spearman’s correlation coefficient, black coefficients show correlation for pooled data, coloured coefficients show cell-type resolved correlations. * p < 0.05, ** p < 0.005, *** p < 0.001, **** p < 0.0001, n.s.: not significant (p > 0.05). Boxes denote interquartile range, horizontal lines are medians. Whiskers show minimum and maximum.

So far, we measured spindle volume as the total volume of all voxels within the segmented spindle mask. Spindle volume, however, might considerably vary from spindle mass (defined as the total steady-state microtubule polymer mass; Reber et al., 2013) depending on microtubule density and spindle architecture. We therefore quantified steady-state spindle mass using Spindle3D in conjunction with the cell volume masks (Figure 5G). For all three cell types examined, spindle mass was directly proportional to spindle volume (r_s_ = 0.94, p = 6 × 10^-88^, Figure S6A) and displayed a comparable scaling relation with cell volume (r_s_ = 0.93, p = 2 × 10^-84^, Figure 5H). Recurrently, spindle width, rather than spindle length, tightly correlated with spindle mass (Figure S6B-C). When evaluating dimensional scaling, spindle length displayed a moderate association with cell volume (r_s_ = 0.75, p = 6 × 10^-35^, Figure 5I), with notably poorer correlation in mESCs (r_s_ = 0.18, p = 0.16), while spindle width robustly scaled with cell volume (r_s_ = 0.80, p = 2 × 10^-44^, Figure 5J) in all three cell lines. Further, we found chromatin volume to linearly scale with cell volume (r_s_ = 0.88, p = 1 × 10^-62^, Figure 5K). Our current understanding of mitotic chromosome scaling with cell volume is limited to so far a single candidate mechanism, i.e. chromatin packing density (as reviewed in Heald and Gibeaux 2018). While studies in *Xenopus* and *C. elegans* show that mitotic chromosome size decreases throughout embryogenesis (Hara et al. 2013; Ladouceur et al. 2015; Kieserman and Heald 2011), systematic and quantitative data from somatic cells are missing. Taken together, our 3D spindle morphometry revealed that spindle volume and mass scale linearly with chromatin volume (but not chromosome number or genome size) and cell volume. Intriguingly, in terms of the spindle’s spatial dimensions, it was not spindle length but rather spindle width that revealed a robust correlation with chromatin and cell volume. Future work shall build on our size scaling analyses to decipher the molecular mechanisms that drive spindle scaling and size control in different species and during development.

## Discussion

So far, early embryonic development, in particular of frogs, fish and worms (Wuhr et al., 2008; Hara and Kimura, 2009; Wilbur and Heald, 2013), has provided experimental models to study spindle scaling and size control. One advantage of early embryonic development is the rapid and dramatic decrease in cell volume over several orders of magnitude. In contrast, somatic cells only show a small variation in cell volume for a given cell type, which makes it harder to discover potential scaling regimes (Marshall 2020). Here, we use volumetric fluorescent microscopy data from somatic cells, stem cells as well as one-cell embryos or zygotes, cells that round up during mitosis or are naturally flat to systematically study scaling relations of spindle, chromatin, and cell geometry. We thereby confirm data from early embryogenesis that within a similar cell size range (V_c_ < 10^6^ µm^3^, d_c_ < 140 µm) spindle volume and spindle mass scale linearly with cell volume. However, whereas it is commonly assumed that spindle length must be tightly tailored to the cell’s dimensions to safeguard fidelity of chromosome segregation and cytokinesis (Goshima and Scholey, 2010), we observed that spindle length was loosely correlated with spindle, chromatin, and cell geometry. Instead, we found the width of the spindle to be a robust predictor of both spindle volume and mass, and concerted with cell and chromatin dimensions.

While it is undebatable that chromatin is sufficient to induce spindle assembly (Heald et al 1996), it remains unclear how its volume, its surface area, its dimensions or a combination of all these factors influence spindle assembly and geometry. In embryonic systems, chromatin content has been shown to only have a minor effect on spindle size. Therefore, it has been suggested that chromatin surface rather than chromatin volume or mass influences spindle size (Brown et al., 2007; Wühr et al. 2008; Dinarina et al., 2009). This is because chromatin triggers spindle self-organization via a diffusion-limited RanGTP gradient, which promotes microtubule nucleation and growth (Gruss et al., 2001). The spatial regulation of microtubule nucleation has recently been shown to determine the upper limit of spindle length (Decker et al., 2018) and particularly in large cells, spindle scaling has been suggested to be governed by microtubule nucleation (Rieckhoff et al., 2020). Chromosomal nucleation, however, might be more relevant in early embryonic systems than in somatic cells (Bird et al., 2008). Furthermore, a combination of modeling and perturbation studies has shown that spindle length is insensitive to the length scale of the Ran gradient in human tissue culture cells (Oh et al. 2016). Thus, how chromatin and the Ran gradient influence microtubule nucleation and dynamics in different scaling regimes remains an exciting open question for future research.

## Material and Methods

### Antibodies

Anti-γ-tubulin (mouse, Sigma T6557), anti-mouse AlexaFluor568 (rabbit, Thermo, A-11061).

### Plasmids

pEGFP-C1-mCherry-CaaX (gift from Paul Markus Müller, Berlin)

### Cell lines

HeLa Kyoto and R1/E mESCs (gifts from Hyman lab, Dresden) and HEK293 (gift from Beckman lab, Berlin), Ptk_2_ cell lines (gift from Simons lab, Dresden).

### Mammalian tissue culture

R1/E mouse embryonic stem cells (mESCs) were cultured in DMEM (high glucose, pyruvate, Gibco) supplemented with 16 % FBS (Gibco), antibiotic-antimycotic (Invitrogen), non-essential amino acids (Gibco), beta-mercaptoethanol (Gibco) and recombinant mouse leukemia inhibitory factor (mLIF, ESGRO). For routine culturing, cells were passaged every 48 hours and seeded at a density of 35,000 cells per cm^2^ onto gelatin-coated dishes. HeLa Kyoto, HEK293 and Ptk_2_ cell lines were cultured in DMEM (high glucose, pyruvate) supplemented with 10 % FBS and antibiotic-antimycotic (Invitrogen) and passaged routinely. Prior to imaging, cells were seeded onto wells of 4-well imaging dishes (Ibidi) or 24-well imaging dishes (Ibidi). To support growth in adherent monolayer, mESCs for seeded onto wells coated with 5 µg/mL laminin-511 (BioLamina) in 1xPBS (supplemented with Ca^2+^, Mg^2+^). All cells were maintained at 37 °C and 5 % CO_2_.

Transfection of HeLa Kyoto cells was performed using Lipofectamine 3000 (Thermo) according to the manufacturer’s instructions.

### Chemical fixation and immunostaining of tissue culture cells

For chemical fixation, R1/E mESCs and HeLa Kyoto cell lines stably expressing tubulin-GFP were seeded 24 h before at a plating density of 100,000 cells per cm^2^, either directly on 24-well imaging slides (Ibidi) or on coverslips. For optimal adherence and monolayer growth, mESCs-designated wells and coverslips were coated with 5 µg/mL laminin 511 in 1x PBS (supplemented with Ca^2+^, Mg^2+^). Media were taken off and replaced with microtubule-optimized fixation buffer (3.2 % paraformaldehyde (EM Sciences), 0.1 % glutaraldehyde (EM Sciences) in 1x BRB80: 80 mM PIPES, 1 mM MgCl_2_, 1 mM EGTA, pH 6.8 with KOH) pre-warmed to 37°C and incubated for 10 min at 37°C. Cells were washed three times in 1x PBS before quenching with 0.1 M glycine (Roth) in 1x PBS for 10 min at RT and 0.1 % NaBH_4_ (Sigma) in 1x PBS for 7 min at RT. DNA was stained with SiR-DNA at a final concentration of 250 nM. Cells were imaged directly in 1x PBS (“fixed only”) or embedded in mounting media (ProLong anti-fade, Invitrogen) and mounted on cover slides (“fixed + mounted”).

For benchmarking of the Spindle3D analysis (see section “Image analysis reliability”, Figure S4A), spindle poles were immunostained using anti-γ-tubulin antibodies (Sigma T6557). Cells grown on 24-well imaging slides (ibidi) were fixed as described above. After quenching, cells were immersed in blocking buffer (3 % BSA, 0.1 % Triton-x 100 in 1x PBS) for 1 h at RT. Primary antibodies were diluted 1:100 in blocking buffer. Incubation with primary antibodies was performed for 1 h at RT under gentle agitation. After three 1x PBS washes for 5 min each, cells were treated with 2 µg/mL anti-mouse AlexaFluor568-labelled secondary antibodies (Thermo, A-11061) in blocking buffer for 45 min at RT and constant agitation. After three final washes with 1x PBS (5 min each), DNA was stained with Hoechst 33343 (Thermo, 62249) at a final concentration of 2 µM and imaged in 1x PBS.

### Image acquisition

For imaging, HeLa Kyoto, Ptk_2_ and HEK293 cells were incubated in imaging medium (FluoroBrite DMEM (Gibco) supplemented with 10 % FBS (Gibco), 4 mM L-glutamine (Invitrogen) and antibiotic-antimycotic (Invitrogen)). mESCs were incubated in stem cell imaging medium (FluoroBrite DMEM (Gibco) supplemented with 16% FBS (Gibco), non-essential amino acids (Gibco), beta-mercaptoethanol (Gibco), sodium pyruvate (Gibco), antibiotic-antimycotic (Invitrogen) and mLIF (ESGRO)). To visualize chromosomes, cells were treated with a final concentration of 250 nM SiR-DNA (Spirochrome). Since Ptk_2_ cells did not show any incorporation of SiR-DNA, we instead incubated the cells for 5 min with Hoechst 33343 (Thermo, 62249) at a final concentration of 2 µM in 1x PBS and replaced the staining solution with imaging medium. Live-cell imaging was carried out using stabilized incubation systems at 37 °C and 5 % CO_2_.

### Imaging was performed on multiple setups

R1/E mESCs were imaged on a Zeiss LSM 800 system (Carl Zeiss Microscopy, Jena, Germany) (sampling in xy: 0.27 µm, z step size: 0.75 µm, total number of slices: 32, pinhole 48.9 µm, unidirectional scan speed: 10, averaging: 2) using a C-Apochromat 40xwater objective (1.2 numerical aperture (NA)), the 488 nm (0.1 % power) and 640 nm (0.1 % power) laser lines and detection ranges of 410 - 558 nm and 586 - 700 nm, respectively. While imaging, cells were incubated using a custom-built incubation chamber (EMBL workshop).

HEK293 cells were imaged on a Nikon spinning disk (CSU-X) confocal system (Nikon Corporation, Tokyo, Japan) equipped with an EMCCD camera (iXon3 DU-888 Ultra, 1024×1024 pixels, 13 μm pixel size) using a 60x Plan Apo oil (1.4 NA) objective (sampling in xy: 0.22 µm, z step size: 0.3 µm, total number of slices: 150), 405 nm (9 % power) and 640 nm (10 % power) laser lines and an excitation time of 200 ms.

Ptk_2_ cells were imaged on the same system using a 40x Plan Fluor oil (1.3 NA) objective (sampling in xy: 0.34 µm, z step size: 0.3 µm, total number of slices: 100 - 150) and 405 nm (10 % power, 100 ms excitation) and 488 nm (18 % power, 300 ms excitation) laser lines.

Again on the same system, HeLa Kyoto cells were imaged using a 60x Plan Apo oil (1.4 NA) objective (see above) or a 100x Plan Apo oil (1.45 NA) objective (sampling in xy: 0.14 µm, z step size: 0.2 µm, z ranges were selected individually per region of interest), using the 488 nm (20 % power, 100 ms excitation) and 640 nm (12 % power, 100 ms excitation) laser lines. Fixed samples of mESCs and HeLa Kyoto cells were recorded on a Nikon spinning disk (CSU-X) confocal system (see above) using a 60x Plan Apo oil (1.4 NA) objective (see above).

Bovine zygotes were generated as previously described (Cavazza et al., 2020). Before fertilization, bovine eggs were injected with 4 pl of mRNAs for mClover3-MAP4-MTBD at 200 ng/µl and of H2B-mScarlet at 60 ng/µl. Bovine zygotes were imaged in 20 µl of BO-IVC (IVF Biosciences) at 38.8°C, 5% CO_2_, 6% O_2_ under paraffin oil in a 35 mm dish with a #1.0 coverslip. Images were acquired with LSM800 confocal laser scanning microscopes (Zeiss) equipped with an environmental incubator box and a 40x C-Apochromat (1.2 NA) water-immersion objective. A volume of 65 µm × 65 µm × 60 µm centered on the chromosomes was typically recorded. The optical slice thickness was 3.00 µm at a z-step size of 2.5 µm. Each zygote was typically imaged every 5 or 10 minutes, using the lowest possible laser intensity (>0.2% for the 488nm laser; >0.2% for the 561 nm laser). mClover3 was excited with a 488 nm laser line and detected at 493 - 571 nm. mScarlet was excited with a 561 nm laser line and detected at 571 - 638 nm.

### Image processing and analysis

After imaging, the only pre-processing step required for downstream analysis is a manual crop of the mitotic cells of interest from the raw files. We suggest using the rectangular selection tool in Fiji (Schindelin et al., 2012). Please note that the morphometric analysis only works on spindles and chromatin that were fully captured in z. Furthermore, the analysis requires fluorescent information of spindle microtubules and chromatin in separate channels that are specified by the user. Other channels will be ignored, but will be displayed in the output image.

The image processing and analysis are implemented in ImgLib2 (Pietzsch et al., 2012). Spindle3D (https://github.com/tischi/spindle3d) is distributed as a Fiji plugin and can be installed by enabling the Spindle3D update site (https://sites.imagej.net/Spindle3D). The image processing and analysis pipeline runs fully automated and consists of the below steps. Parameters are shown in quotation marks, with the default parameter values indicated after a colon.

#### Isotropic resampling

To facilitate implementation of the image analysis algorithms all channels of the input image are resampled to an isotropic voxel size (“voxel size for analysis”: 0.25 µm).

#### Metaphase plate initial segmentation

In order to find the location of the metaphase plate in the image, we rely on the fact that the DNA signal in condensed chromosomes is brighter than interphase DNA. To find an intensity threshold above which voxels belong to condensed (metaphase) chromosomes, we first downsample the DNA image such that the width of one voxel resembles the typical width of the metaphase plate (“voxel size for initial DNA threshold”: 1.5 µm) (see Figure S1A, left). The intensity values in the downsampled image are computed by averaging with a gaussian blur with a sigma of half the “voxel size for initial DNA threshold”. In this downsampled image we find the maximal (max) and minimal (min) intensity. We empirically determined that (max+min)/2 serves as a reliable threshold. We then apply this threshold to the DNA image to create a binary mask and perform a connected component labelling (Figure S1A, center). We remove all connected components that touch the lateral (xy) image boundary and of the remaining ones only keep the largest one, which we define to be the initial metaphase plate object.

#### Metaphase plate center and orientation

To determine the orientation of the metaphase plate, we use an algorithm from ImageJ’s 3D Image Suite (Ollion et al., 2013) to fit a 3D ellipsoid to the initial metaphase plate object, resulting in three vectors along the shortest, middle, and longest axes as well as the coordinates of the metaphase plate center. To facilitate the implementation of the subsequent algorithms, e.g. in terms of specifying ranges to be included in certain computations, and also to facilitate visual inspection of the images, we use these vectors to compute a transformation that puts all images into a new coordinate system such that the new z-axis corresponds to the shortest axis of the metaphase plate, roughly corresponding to the spindle pole-to-pole axis, and such that the origin of the coordinate system coincides with the center of the metaphase plate. We will refer to these transformed images as metaphase plate aligned images (Figure S1A, right).

#### Metaphase plate width

Using the metaphase plate aligned DNA image, we compute an average intensity profile along the shortest DNA axis (z-axis of the aligned image), limiting the computations to a maximum width that is based on the extent of the shortest axis of the initial ellipsoid fit times 2. We then compute the derivative of this profile at a resolution of “metaphase plate derivative delta”: 3 µm. We define the metaphase plate width as the distance between the locations with the highest absolute values in the derivative (see Figure S1B, left). This procedure is motivated by the fact that, due to the various chromosomes and the diffraction limit of the microscope, the metaphase plate has an overall irregular appearance. For example, our analysis is robust to an individual chromosome “sticking out” of the metaphase plate as this will not shift above maxima of the derivatives (see Figure S1B, left). In other words, our approach measures an average width, determined by the average position of all chromosomes.

#### Metaphase plate length

Using the metaphase plate aligned DNA image, we compute an average lateral radial intensity profile (see Figure S1B, center left), limiting the computations to a maximum length determined by the extent of the longest axis in the initial ellipsoid fit times 2. We define the metaphase plate length as 2 times the distance between the origin to the position of the minimum in the derivative of the intensity profile. Again, this approach reports an average measurement that is robust to any details that the various arrangements of the chromosomes in the metaphase plate may have. In addition, both the measurements of the metaphase width and length have the advantage of not depending on the choice of any intensity threshold.

#### Chromatin dilation

We further utilize the average lateral radial DNA intensity profile (s.a.) to calculate the ratio of the intensity in the center of the metaphase plate (position zero along the radial profile) and the brightest part along the profile (see Figure S1B, center right). This ratio is subtracted from 1 to report on the magnitude of the dilation in the center of the metaphase plate, higher values corresponding to a more pronounced opening, while small values reflect homogeneously closed metaphase plates.

#### Chromatin volume

In order to facilitate the comparison with previously published measurements, we decided to adopt the method published in Hériché et al., (2014), where the Otsu algorithm (Otsu, 1979) is used to determine an intensity threshold and the chromatin volume is determined as the sum of the volume of all voxels above this threshold. The Otsu algorithm relies on a bimodal (foreground and background) intensity distribution. We therefore apply the Otsu algorithm to a region of interest determined by the previously measured metaphase plate width and length (see Figure S1B, right) where the intensity values only comprise the metaphase plate (foreground) and parts of the cell devoid of DNA signal (background), but exclude (unwanted) DNA signal from surrounding cells. We apply the determined threshold to the whole image, perform a connected component analysis, remove regions touching the image borders and keep the largest region, which we call segmented metaphase plate. The volume of this region is the chromatin volume.

#### Spindle segmentation

We explored various methods of reliably determining an intensity threshold for assigning pixels to the mitotic spindle and developed an algorithm relying on the observation that, in all data we analysed, the metaphase plate length was always substantially (on average 25%) larger than the spindle width. The direct vicinity of the metaphase plate therefore contains a substantial fraction of pixels inside as well as outside of the spindle. Thus, this region is well suited for determining an automated threshold using the Otsu algorithm (see section “Chromatin volume”). Technically, we apply the Otsu algorithm to all tubulin intensity values in a rim of a thickness of 1 pixel around the segmented metaphase plate (see Figure S1C, center left). We then apply this threshold to the whole tubulin image and perform a connected component analysis. As there can be other cells with relatively bright tubulin intensities in the same image, we filter the regions, only keeping regions where at least one of their pixels is within a defined distance to the center of the metaphase plate (“spindle fragment inclusion zone”: 3 µm). We will refer to the union of those regions as the spindle mask.

#### Spindle volume

The spindle volume is computed as the volume of all voxels in the spindle mask (see Figure S1C, center left).

#### Spindle average intensity

The average gray value of all tubulin voxels within the spindle volume mask.

#### Spindle intensity variation

Spindles have different degrees of homogeneity in terms of their distribution of polymerised tubulin. We measure this by computing the coefficient of variation of the (threshold subtracted) tubulin intensities within the spindle mask.

#### Spindle poles locations

In our algorithm, both spindle length and spindle orientation are determined by the vector that connects the two spindle poles. We locate the spindle poles in two steps. First, we draw a line profile through the spindle mask along the shortest metaphase plate axis and through the metaphase plate center. The two locations along the line profile where the spindle mask intensity drops from 1 to 0 (i.e. the spindle mask ends) are the two initial spindle poles (see Figure S1C, center right). As the spindle axis is often not completely aligned with the shortest metaphase plate axis, the initial spindle poles need to be refined. To do so, we determine the locations of the pixels with the maximum intensity in the tubulin image in a small neighborhood around the initial spindle poles. The extent of this neighborhood is controlled by the “axial pole refinement radius”: 1.0 µm and the “lateral pole refinement radius”: 2.0 µm, where “axial” refers to along the shortest metaphase plate axis and “lateral” refers to the perpendicular directions (see Figure S1C, center right).

#### Spindle center location

We define the middle between the two spindle poles as the spindle center location.

#### Spindle center to metaphase plate center distance

The distance of the metaphase plate center (s.a.) to the spindle center.

#### Spindle length

We define the spindle length as the distance between the two spindle poles (see Figure S1C, center right).

#### Spindle angle

The two spindle poles allow to define a spindle axis vector that points from one pole to the other. We apply below formula (in the coordinate system of the input data where we assume the coverslip plane to be perpendicular to the z-axis) to compute the angle of the spindle axis and the coverslip plane as follows: 90.0 - abs(angle_degrees(z-axis, spindle-axis)). For the computation of an angle between two axes there are always two solutions. Here, the computations within the function angle_degrees are done such that the smaller one, i.e. with a value between 0 and 90 degrees, is picked (see Figure S1C, right).

#### Spindle coordinate system

We define a new coordinate system in which the spindle center is at the origin and the spindle poles are aligned along the z-axis. This coordinate system simplifies the following measurement.

#### Spindle widths

To measure the spindle width we perform a maximum projection of the spindle mask along the spindle axis (the z-axis in the spindle coordinate system). We smoothen the 2D projected spindle mask by a morphological opening operation with a radius of 2 pixels. We then compute the width of the binary mask in steps of 10 degrees (see Figure S1C, center right). We define the mean of the resulting widths as the “average spindle width”. To capture potential anisotropies in the spindle shape, we fetch both the minimum and maximum of the width at all measured angles, resulting in the outputs: “minimal spindle width” and “maximal spindle width”.

#### Spindle aspect ratio

As a measure for spindle shape, we define the ratio of spindle length and the average spindle width as the spindle aspect ratio.

#### Tabular output

The plugin outputs all measured values in a table, the column names corresponding to the respective measurements.

#### Image output

The plugin also outputs a multichannel image, composed of the DNA and tubulin signal, the DNA mask, the spindle mask, and another image containing three points corresponding to the spindle poles and the spindle centre (see Figure 2B). All images are sampled isotropically at the voxel size for analysis. For ease of inspection, all images are aligned such that the x-axis corresponds to the measured spindle axis and such that the center of the image corresponds to the spindle center.

### Cell volume quantification

Cells expressing tubulin genetically fused to a fluorescent protein show characteristic cytoplasmic background fluorescence (see Figure 5A). We used the machine-learning based segmentation software Ilastik (Berg et al., 2019) to train pixel-classification models to distinguish between true mitotic cytoplasm and all other voxels. Before training and prediction, all images were rescaled to an isotropic voxel size of 0.25 µm to be consistent with the analysis voxel size applied in the Spindle3D analyses. For training, training set images were annotated in the auto-context module in Ilastik, a two-step workflow, where the second stage receives the prediction results from the initial stage. In the first stage, the default random forest algorithm was trained with the classes “spindle microtubules”, “mitotic cytoplasm”, “interphase microtubules”, and “background”. Brightness features were excluded, to avoid bias in fluorescence signal strength, i.e. varying expression levels of fluorescent tubulin. We used all available texture and edge filters for training. In the second stage, we trained another random forest to distinguish between the two classes “true mitotic” and “other”, while again all brightness features were ignored and all texture and edge features were included. After successful training, batch processing was performed using the ilastik integration in Fiji. The second-stage probability masks (see Figure 5A, center) were de-noised with a 3D Gaussian filter (sigma = 2 pixels) and thresholded at the cutoff value 0.5, reflecting the binary prediction approach to distinguish between “true mitotic” and “other”. The volumes of the resulting segmentation masks (see Figure 5A, right) were quantified using the “3D analyse regions” function in the MorphoLibJ package (Legland et al., 2016).

### Spindle mass quantification

In cells expressing fluorescently-tagged tubulin, we can define the average voxel gray value within the cell mask (see section “Cell volume quantification”) as the average concentration of tubulin in the whole cell [T]_C_. Analogously, the average concentration of tubulin in the spindle [T]_S_ is reflected by the average voxel value within the spindle mask. To account for system-internal noise of the imaging setup, we calculated the median voxel values in the bottommost slices of the image stack. We then subtracted this value from both [T]_C_ and [T]_S_. We define spindle mass as the sum of tubulin (free and polymer) within the spindle volume V_S_. To correct for cell-specific tubulin-GFP expression levels, we normalised spindle masses by the cell-specifc fluorescence correction factor Fl_corr_ ∼ [T]_C_: Spindle mass = [T]_S_ * V_S_ / Fl_corr_.

### Image analysis reliability and accuracy

In order to benchmark the measurements of the plug-in, we labelled centrosomes of tubulin-GFP expressing R1/E mouse embryonic stem cells with anti-γ-tubulin antibodies (see section “Chemical fixation and immunostaining of tissue culture cells”). Having imaged the spindles using confocal microscopy, in Fiji we located the outward-facing edges of the γ-tubulin signals and defined their 3D coordinates as the ground-truth spindle poles (see Figure S6A). The euclidean distance between the two poles was defined as the ground-truth spindle length. In parallel, we located the 3D coordinates of the poles exclusively by looking at the tubulin-GFP signals. The euclidean distance between this pair of poles was defined as the manual spindle length. Ultimately, we compared both the ground-truth spindle length and the manual spindle length measurements to measurements derived via the Spindle3D plug-in.

Opposed to spindle length, we lack proper ground-truth references for spindle width. Nevertheless, we benchmarked the plug-in’s performance against human measurement. To this end, we made use of the spindle axis registration of the Spindle3D analysis and verified the orientation with the centrosomal γ-tubulin signals (see above). In Fiji, we used Image > Stack > Reslice to match the direction of the spindle axis with the z axis and performed a maximum projection. Using only the tubulin-GFP and γ-tubulin signals, we manually determined 4 extents of the spindle in the projected image (see Figure S6B) and calculated their mean to serve as the manual spindle width reference measurement.

To verify the performance of the pixel classification-based mitotic cell volume quantification, we transfected tubulin-GFP expressing HeLa Kyoto cells with plasmids encoding mCherry tagged with the CaaX motif (Clarke, 1992) for cell membrane localisation. We acquired confocal images of mCherry-positive mitotic cells (see Figure S6E) and performed our tubulin-GFP based cell volume quantification as described above. In parallel, we manually segmented cell volumes using the mCherry-CaaX landmark channel and the volume manager in the SCF MPI-CBG Fiji package (https://sites.imagej.net/SCF-MPI-CBG/) (see Figure S6F) to generate a binary cell mask, the volume of which was quantified via the “3D analyse regions” function in the MorphoLibJ package (Legland et al., 2016).

For quality control, 3D image stacks were rendered using the multichannel visualisation package ClearVolume (Royer et al., 2015) in Fiji.

### Data analysis and visualization

After image analysis, we used the pandas (The pandas development team, 2020), SciPy (Virtanen et al., 2020) and NumPy (Harris et al., 2020) libraries in Python to further analyse the data. Statistical tests (Wilcoxon signed-rank test) were carried out using the scipy.stats package. ANOVA and post-hoc testing was performed using the Python statsmodel package (Seabold et al., 2010). Power-law fitting was performed in the scipy.optimize library. Linear regression was performed in the scikit-learn package (Pedregosa et al., 2011). All data visualization was carried out in the Python Altair library (VanderPlas et al., 2018) and assembled in Adobe Illustrator 2021 (Adobe inc. 2020).

## Acknowledgments

We thank all present and past members of the Reber lab and the Advanced Light Microscopy Facility (ALMF) at the European Molecular Biology Laboratory (EMBL) for support, in particular Aliaksandr Halavatyi, Faba Neumann, and Stefan Terjung. We thank Jan Schmoranzer from the Advanced Medical BIOimaging Core Facility, Charité Berlin for imaging support. We are grateful to Jean-Karim Hériché and Julius Hossain (both EMBL) for advice on image analysis. We thank Renata Basto and Melina Schuh and their labs for testing the plug-in and for critical discussions. We are grateful to Tommaso Cavazza (Schuh lab) for kindly providing the bovine zygote dataset. We thank Christopher Schmied and Jan Brugues for critical comments on the manuscript.

This work was supported by the Deutsche Forschungsgemeinschaft (DFG, A Quantitative Force Map of the Mitotic Spindle, RE 3925/1-1 to S.R.) and by iNEXT (grant number 653706, project PID 3503 funded by the Horizon 2020 program of the European Union to S.R.). S.R. further acknowledges funding by the IRI Life Sciences (Humboldt-Universität zu Berlin, Excellence Initiative/DFG).

## Author contributions

Conceptualization, S. Reber; funding acquisition, S. Reber; Investigation, T. Kletter carried out all experiments. T. Kletter and C. Tischer constructed the plugin, analyzed images, and performed data analysis; Writing–Original Draft, S. Reber with input from T. Kletter and C. Tischer. S. Reusch and N. Dempewolf acquired selected datasets and performed quality control analyses.

---

**Figure S1:**
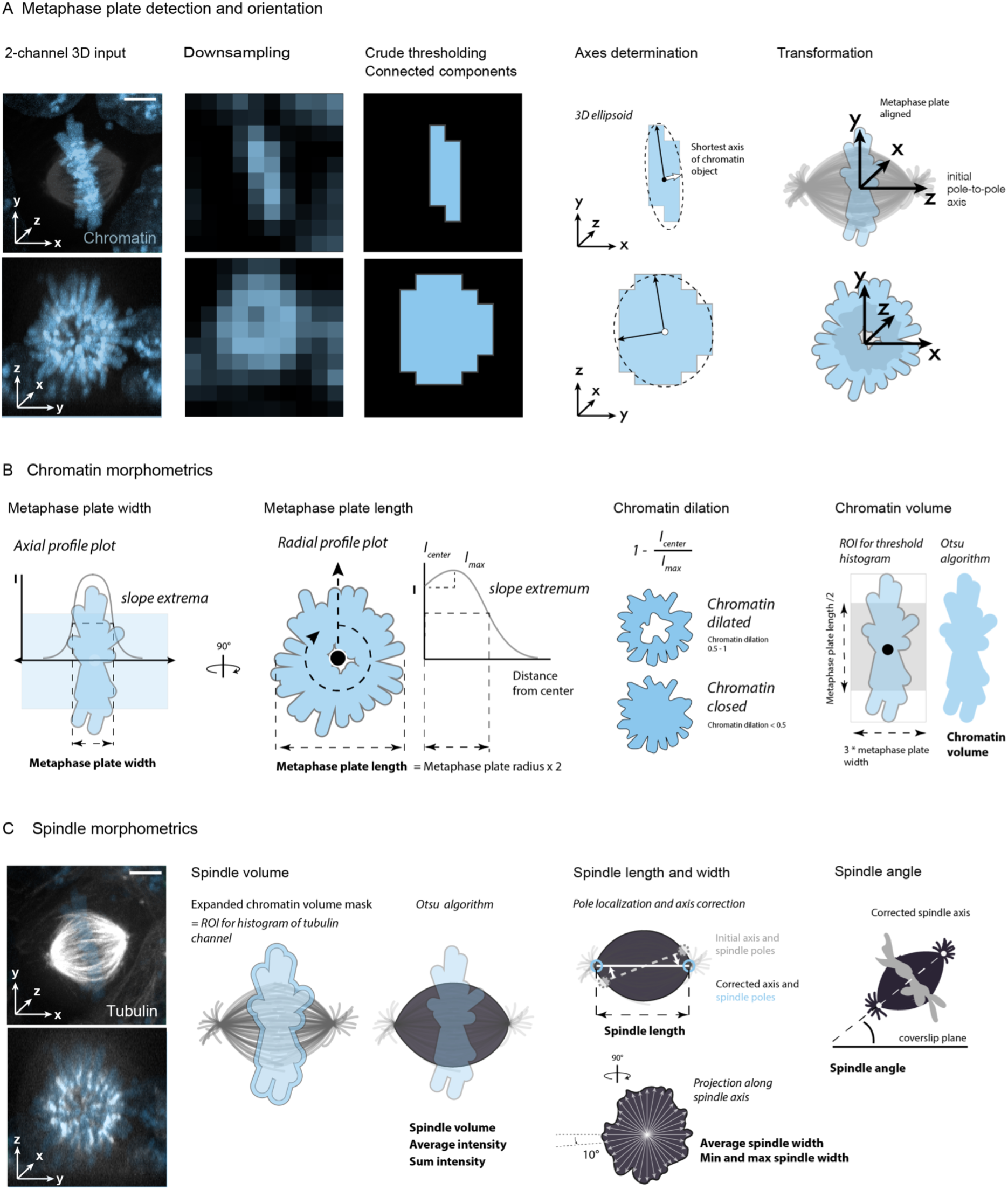
Spindle3D morphometric analysis workflow. **(A)** In confocal micrographs, mitotic cells (chromatin shown in blue, tubulin in grayscale) are detected by crude, histogram-based segmentation and connected-component analysis. The shortest axis of the metaphase plate object is determined and initiates an imaging axis-independent coordinate system. **(B)** In the newly aligned image, radial and axial intensity profiles serve as robust guides to quantify the extents of the usually irregularly shaped chromatin plate. Moreover, radial profiles inform on the magnitude of dilation of the metaphase plate. Based on the extents of the plate, a three-dimensional region of interest is used to limit the pixels considered for histogram-based segmentation of the chromosomes, excluding potentially interfering signals from nuclei in close proximity. **(C)** Analogously, only a fraction of the tubulin channel pixels (the ones immediately bordering the chromatin mask and thus either represent spindle microtubules, or non-spindle tubulin inside the cell) are considered for Otsu thresholding the spindle. Spindle poles are the brightest pixels found within defined radii around the intersections of the initial spindle axis (found in (A)) and the spindle volume mask. This mask is projected along the now corrected spindle axis. The resulting area is radially scanned in 10° steps, to ultimately measure 18 lateral spindle extents, their mean representing the average spindle width. Finally, the tilt of the corrected spindle axis is used to determine the spindle angle.

**Figure S2:**
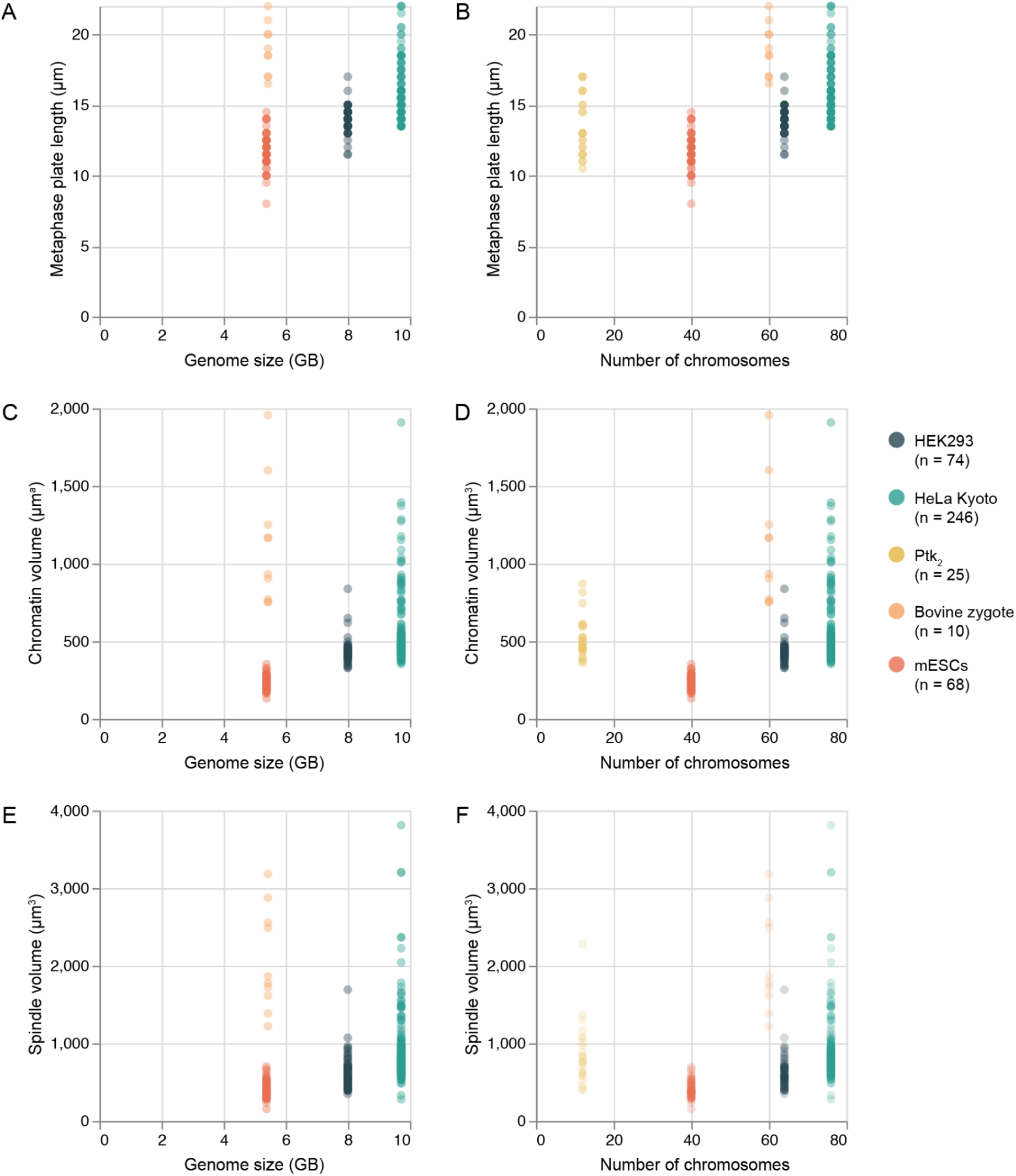
Relationship between genome sizes and spindle and chromatin dimensions. HEK293 cells were described as hypo-triploid (3n-) (Bylund et al., 2004) with an average chromosome number of 64. Considering the median human diploid genome size of 5.72 gigabases (Gb) (NCBI) and the diploid human chromosome number of 46, we estimated the genome size to be approximately 8.00 Gb. HeLa cells were described as hyper-triploid (3n+) (Macville et al., 1999) with an average chromosome number of 76, we estimated the average HeLa genome to amount to approx. 9.72 Gb. The diploid genome of *Mus musculus* corresponds to 40 chromosomes. The median diploid genome size is reported to be 5.38 Gb (NCBI). Analogously, the diploid genome of *Bos taurus* corresponds to 60 chromosomes, the median diploid genome size is 5.44 Gb (NCBI). The genome of the marsupial species *Potorous tridactylus* is not yet sequenced, the female diploid chromosome number is 12 (Rens et al., 1999). **(A)** Scatter plot displaying the relationship between genome size and metaphase plate length and **(B)** number of mitotic chromosomes and metaphase plate length. **(C)** Scatter plot displaying the relationship between genome size and chromatin volume and **(B)** number of mitotic chromosomes and chromatin volume. **(E)** Scatter plot displaying the relationship between genome size and spindle volume and **(F)** number of mitotic chromosomes and spindle volume. Circles represent single cells.

**Figure S3:**
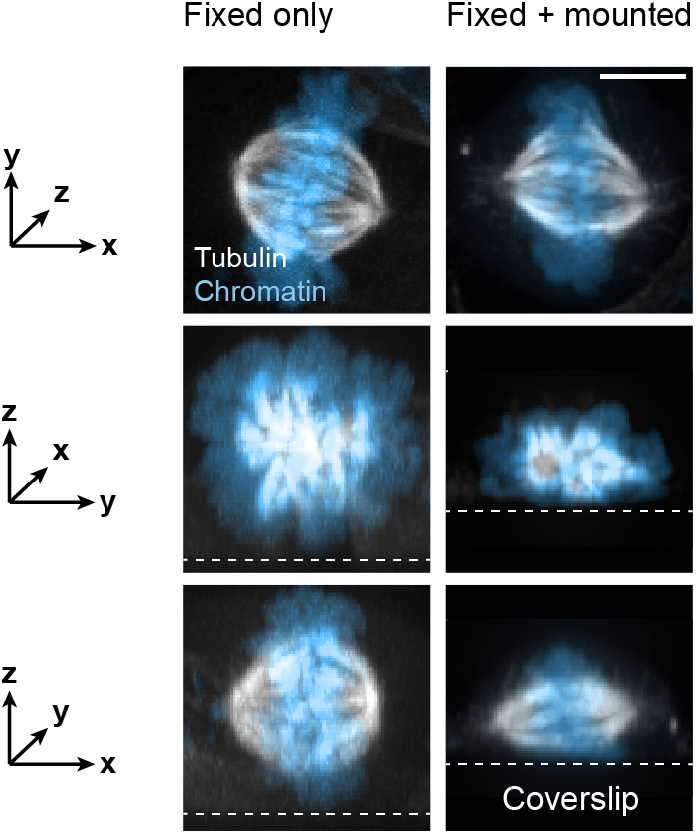
Sample handling distorts spindle shape. Left column shows projections along the imaging axis (top), along the x axis (middle) and along the y axis (bottom) of a chemically fixed mouse embryonic stem cell (mESC) imaged in 1x PBS. Analogously, the right column shows a mESC that, after fixation, was embedded in mounting solution and mounted on a cover slide. Tubulin-GFP is shown in grayscale, Hoechst-labelled chromatin in blue. The dotted lines indicate the plane of the cover glass. Scale bar: 5 µm.

**Figure S4:**
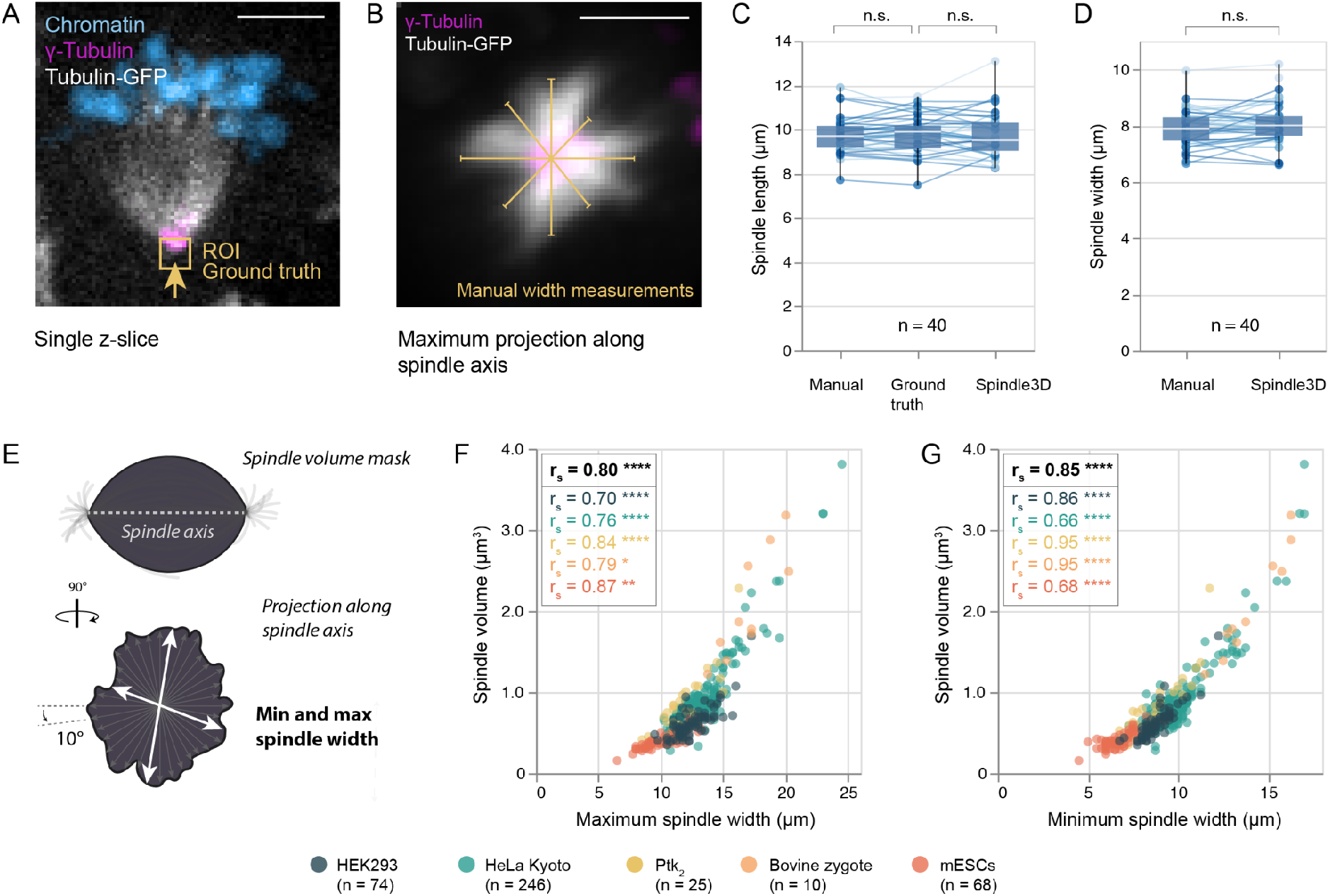
Performance accuracy of manual versus automated spindle measurements. **(A)** Micrograph (single z-slice) of a tubulin-GFP (gray scale) expressing mouse embryonic stem cell (mESC) fixed at mitosis. Antibody stainings were used to detect γ-tubulin (magenta). DNA is stained with Hoechst (blue). The arrow and region of interest (ROI, yellow) highlight the outer edge of the γ-tubulin signal, the position considered as ground truth spindle pole. Scale bar: 5 µm. **(B)** Maximum projection along the spindle axis of a tubulin GFP-expressing mitotic mESC with labelled centrosomes (γ-tubulin, magenta). Four manually drawn spindle width measurements (yellow) were averaged to yield the reference spindle width. Scale bar: 5 µm. **(C)** Box plots show distributions of spindle length measurements (n = 40) derived by manually placing spindle poles within the tubulin-only 3D image (“Manual”), or by manually placing spindle poles within the γ-tubulin-only 3D image (“Ground truth”) or by subjecting the chromatin/tubulin stack to analysis by Spindle3D. **(D)** Box plots show distributions of spindle width measurements (n = 40) derived manually or via Spindle3D. Boxes reflect the interquartile range, whiskers show the minimum and maximum. The medians are shown as horizontal white lines inside the boxes. Circles reflect measurements on individual spindles and are linked across the methods by lines. Hypothesis testing was performed using the Wilcoxon signed-rank test. N.s: not significant (p > 0.05). **(E)** Rationale for determining the minimum and maximum spindle width after segmentation in Spindle3D. **(F)** Scatter plots showing the relationship between the maximum spindle width and spindle volume, **(G)** between the minimum spindle width and spindle volume. Circles represent individual cells. r_s_: Spearman correlation coefficient. * p < 0.05, ** p < 0.005, *** p < 0.001, **** p < 0.0001, n.s.: not significant (p > 0.05).

**Figure S5:**
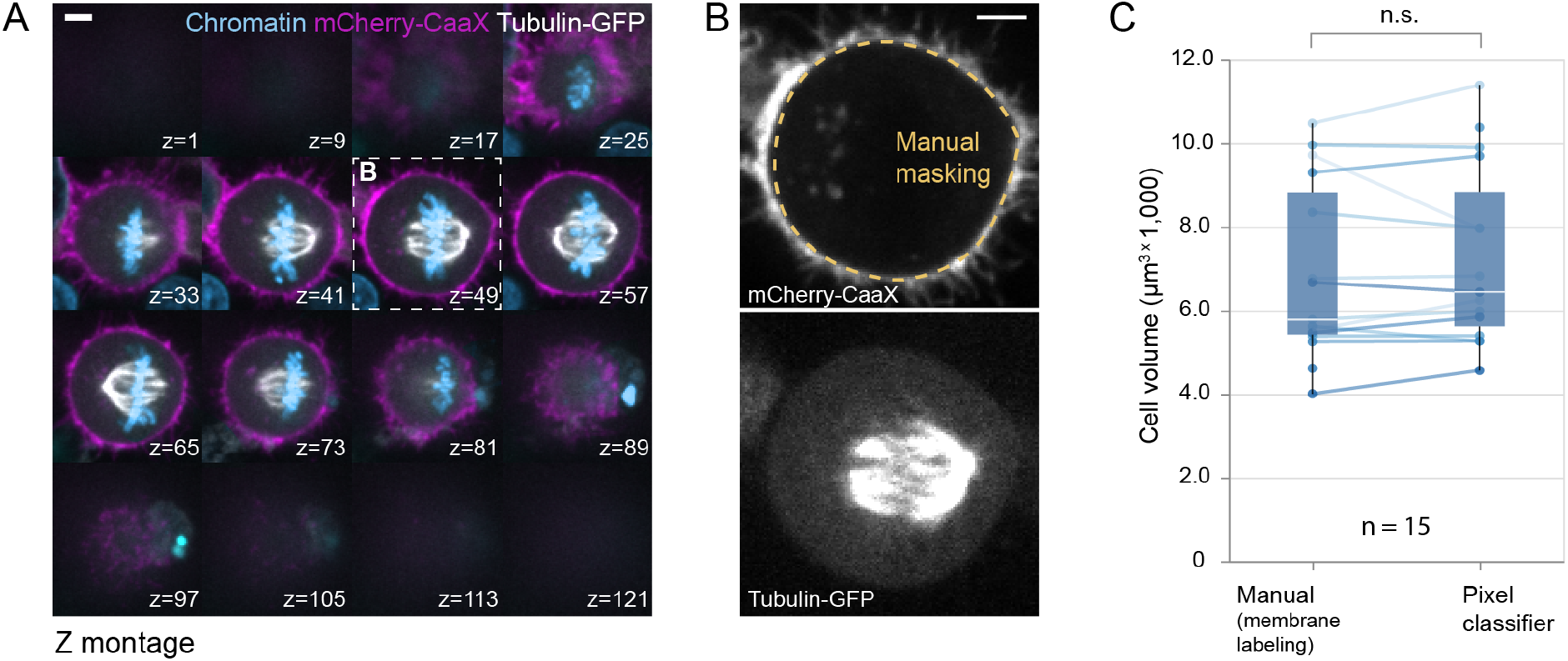
Performance accuracy of manual versus automated cell volume measurements. **(A)** Z-montage showing a mitotic HeLa Kyoto cell expressing tubulin-GFP and mCherry-CaaX. Scale bar: 5 µm. **(B)** Isolated slice of (A) highlighting the cell membrane landmark channel (top) with the manually traced cell boundary (yellow) and the tubulin-GFP channel (bottom) used in pixel classification-based segmentation. Scale bar: 5 µm. **(C)** Distributions showing cell volumes (n = 15) as determined in the landmark channels versus through pixel classification in the fluorescent tubulin channel. Boxes reflect the interquartile range, whiskers show the minimum and maximum. The medians are shown as horizontal white lines inside the boxes. Circles reflect measurements on individual spindles and are linked across the methods by lines. Hypothesis testing was performed using the Wilcoxon signed-rank test. N.s: not significant (p > 0.05).

**Figure S6:**
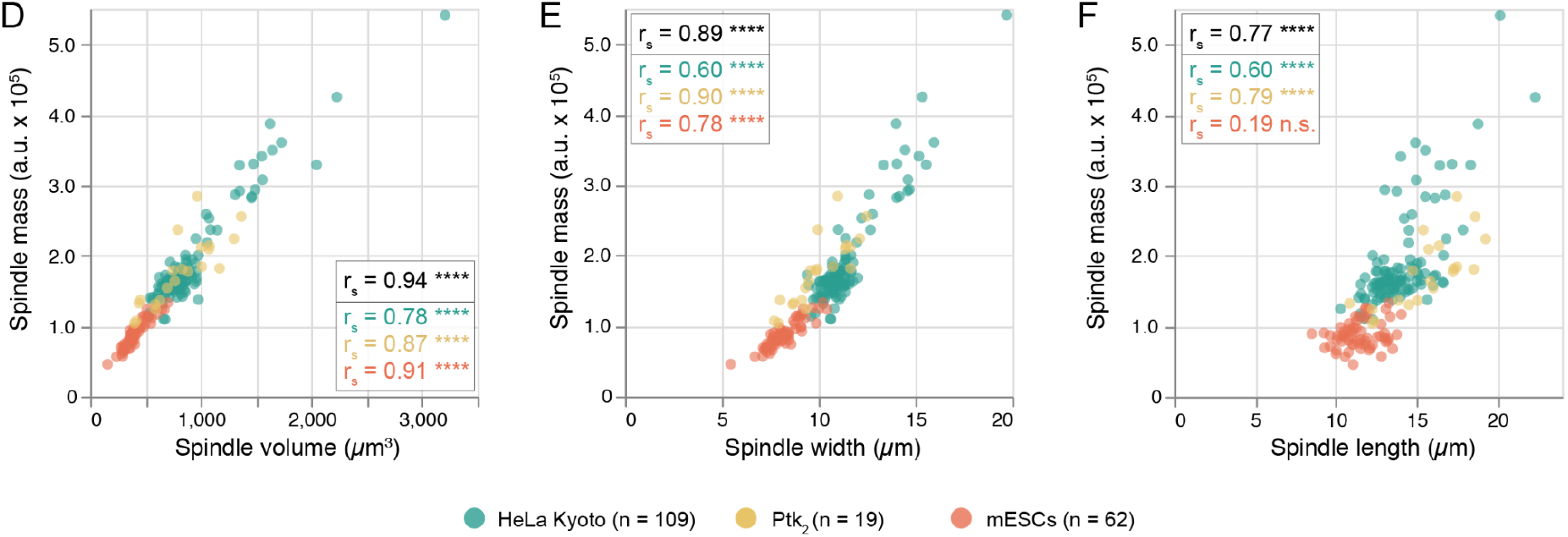
Spindle mass is directly proportional to spindle volume. **(A)** Scatter plot displaying the relationship between spindle volume and spindle mass. **(B)** The relationship between spindle width and spindle mass and **(C)** the relationship between spindle length and spindle mass. Circles represent individual cells. r_s_: Spearman correlation coefficient. * p < 0.05, ** p < 0.005, *** p < 0.001, **** p < 0.0001, n.s.: not significant (p > 0.05).

